# Induction of the inflammasome by the SARS-CoV-2 accessory protein ORF9b, abrogated by small-molecule ORF9b homodimerization inhibitors

**DOI:** 10.1101/2024.05.31.596900

**Authors:** Erika Zodda, Mònica Pons, Natàlia DeMoya-Valenzuela, Cristina Calvo-González, Cristina Benítez-Rodríguez, Blanca Díes López-Ayllón, Achraf Hibot, Marta Cascante, María Montoya, María Dolors Pujol, Jaime Rubio-Martínez, Timothy M. Thomson

**Author notes:** Correspondence: Timothy M Thomson, IBMB-CSIC, CIBER-EHD and Universidad Peruana Cayetano Heredia,; Jaime Rubio Martínez, Universitat de Barcelona.

## Abstract

Viral accessory proteins play critical roles in viral escape form host innate immune responses and in viral inflammatory pathogenesis. Here we show that the SARS-CoV-2 accessory protein, ORF9b, but not other SARS-CoV-2 accessory proteins (ORF3a, ORF3b, ORF6, ORF7, ORF8, ORF9c, ORF10), strongly activates inflammasome-dependent caspase-1 in A549 lung carcinoma cells and THP-1 monocyte-macrophage cells. Exposure to lipopolysaccharide (LPS) and ATP additively enhanced the activation of caspase-1 by ORF9b, suggesting that ORF9b and LPS follow parallel pathways in the activation of the inflammasome and caspase-1. Following rational *in silico* approaches, we have designed small molecules capable of inhibiting the homodimerization of ORF9b, which experimentally inhibited ORF9b-ORF9b homotypic interactions, caused mitochondrial eviction of ORF9b, inhibited ORF9b-induced activation of caspase-1 in A549 and THP-1 cells, cytokine release in THP-1 cells, and restored type I interferon (IFN-I) signaling suppressed by ORF9b in both cell models. These small molecules are first-in-class compounds targeting a viral accessory protein critical for viral-induced exacerbated inflammation and escape from innate immune responses, with the potential of mitigating the severe immunopathogenic damage induced by highly pathogenic coronaviruses and restoring antiviral innate immune responses curtailed by viral infection.

## Introduction

Coronaviruses are unique among Nidoviruses in that they encode a variable number of accessory proteins whose functions are not essential for virus replication, yet play a relevant role in pathogenesis ^1^. SARS-CoV-2, the causative agent of COVID-19, is an enveloped virus consisting of a positive-sense, single-stranded RNA genome ^2^. Two overlapping open-reading frames, ORF1a and ORF1b, are translated from the positive-strand genomic RNA and generate continuous polypeptides, which are cleaved into a total of 16 nonstructural proteins (NSPs). The remaining genomic regions encode four structural proteins: spike (S), envelope (E), membrane (M), and nucleocapsid (N), and six annotated accessory proteins (ORF3a, 6, 7a, 7b, 8, and 10). SARS-CoV-2 also harbors several unannotated accessory ORFs, including alternative open reading frames within ORFs S (ORF2d), N (ORF9b, ORF9c), and ORF3a (ORF3b, ORF3c, ORF3d) ^3^.

Host-virus interactome analyses have uncovered human proteins that physically associate with SARS-CoV-2 proteins and that may participate in the virus life cycle, infection, replication, and budding ^4–9^. Among these, interactions with mitochondrial proteins are particularly abundant. The interaction of SARS-CoV-2 proteins with mitochondrial proteins results in mitochondrial dysfunction and increased oxidative stress, ultimately leading to the loss of mitochondrial integrity and cell death ^10–12^. Recent studies suggest that the involvement of mitochondria in SARS-CoV-2 infection is a hallmark of disease pathology ^13–15^ and that it may also explain relevant disease patterns in post-acute COVID syndromes ^15,16^.

As a result of compromised mitochondrial integrity, oxidized mitochondrial DNA, cardiolipin and cytochrome c are released into the cytosol, activating damage-associated molecular pattern (DAMP) receptors and Nucleotide-binding oligomerization domain, Leucine rich Repeat and Pyrin domain (NLR)-containing proteins, including NLRP3, with consequent hyper-activation of the inflammasome, leading to sustained local and systemic inflammatory responses ^17^. Like other viruses, SARS-CoV-2 can activate the inflammasome, thus contributing to unmitigated activation of the innate immune system leading to pathogenicity ^18–23^. Several SARS-CoV-2 structural proteins, including spike (S), nucleocapsid (N), envelope (E) or non-structural proteins, such as nsp6 or ORF3a, have been reported to induce the activation of the inflammasome ^24^. Whether these diverse proteins share common mechanisms to activate the inflammasome is not currently known.

The SARS-CoV-2 accessory protein ORF9b is a small 97 amino acid polypeptide that, in infected cells, forms homodimers that associate with lipids and cell membranes, or monomers that interact with the mitochondrial translocase, TOM70 ^5,25–29^. The interaction of an ORF9b monomer with TOM70 competes with the binding of the chaperone HSP90 to TOM70, thus leading to its diminished stability and function as a mitochondrial translocase ^26,27,30^ and also its ability to interact with MAVS ^28^(28) and other critical components of the signaling pathways that induce type I interferons in response to cytosolic viral RNAs ^28,31,32^. As such, ORF9b is a major factor used by SARS-CoV-2 to interfere with the activation of innate immunity upon viral infection of cells and thus to favor unimpeded viral replication and spread. It follows that inhibition of these ORF9b activities would very likely favor more effective defenses against SARS-CoV-2 infection, lowering viral pathogenesis. Importantly, ORF9b is conserved among SARS-CoV-2, SARS-CoV and other coronaviruses both in sequence and function ^4,5,26,29,33,34^. To date, no small-molecule ORF9b inhibitors have been reported.

Here, we show that ORF9b induces the inflammasome-dependent activation of caspase-1, without requiring a priming event. This induction is nearly one order of magnitude more potent than that of other SARS-CoV-2 accessory proteins and follows a pathway distinct from that of exogenous lipopolysaccharide (LPS) and ATP. Through virtual design and screening, we have developed novel small molecule inhibitors of ORF9b homodimerization that exert a potent antagonism of the pro-inflammatory activities of ORF9b and restore the type I interferon responses antagonized by ORF9b.

## Results

### SARS-CoV-2 ORF9b potently induces caspase-1 activation

In order to determine the ability of SARS-CoV-2 accessory proteins to modulate the activation of the inflammasome, we assessed caspase-1 activity, as a proxy of inflammasome activation, of A549 lung carcinoma cells either transiently transduced with, or stably integrating, ORF3a, ORF3b, ORF6, ORF7a, ORF7b, ORF8, ORF9b, ORF9c or ORF10 under basal conditions or after exposure to the toll-lie receptor 4 (TLR4) agonist, lipopolysaccharide (LPS) followed by extracellular ATP (LPS+ATP). Both stable, transiently transduced and long-term constitutive expression of SARS-CoV-2 accessory proteins impacted caspase-1 activation to similar degrees, perhaps with the exception of ORF7b (Fig. 1A,B). Uniquely, expression of ORF9b strongly induced caspase-1 activity under both experimental conditions and in the absence of priming events (Fig. 1A,B). On the other hand, both transient and long-term expression of ORF7a, but only transient expression of ORF3d, suppressed basal caspase-1 activity (Fig. 1,B). As expected, exposure to LPS+ATP also strongly induced caspase-1 activity in control A549.pLex cells (Fig. 1A,B) and it exerted an additive effect on the activation of caspase-1 induced by ORF9b (Fig. 1A,B), which suggests that ORF9b and LPS+ATP may use distinct pathways to activate caspase-1. Of note, both transient and long-term expression of ORF3a, ORF7a, ORF8 and ORF10, as well as long-term expression of ORF7b, inhibited the activation of caspase-1 induced by LPS+ATP in control A549.pLex cells (Fig. 1A,B).

**Figure 1.**
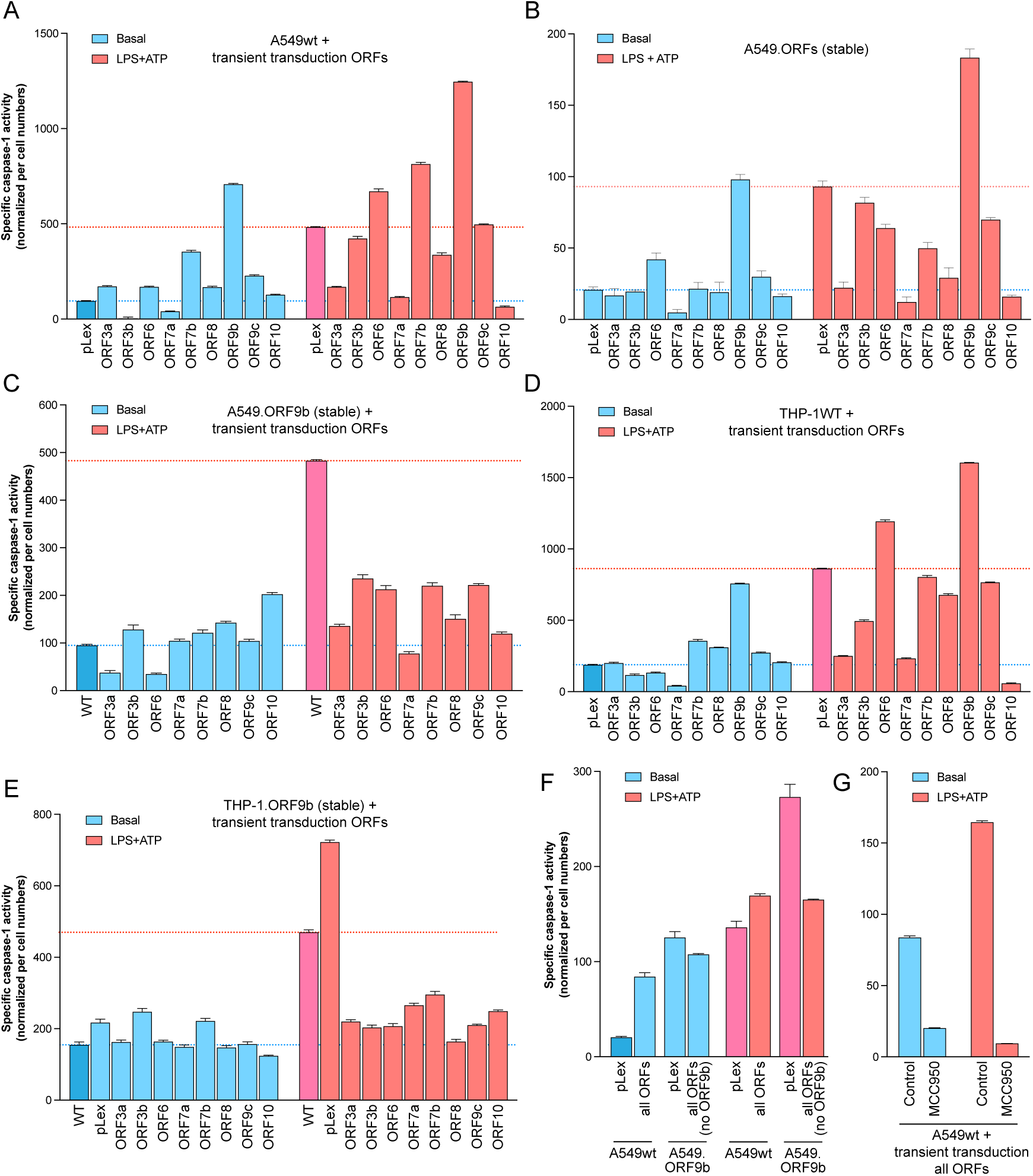
Impact of SARS-CoV-2 accessory proteins on caspase-1 activity. **A**. Transient expression of ORF9b strongly induces caspase-1 activity in A549 cells. Parental A549 cells (A549wt) were transduced with lentiviral particles for the expression of the indicated accessory proteins and released caspase-1 activity determined 48 h after transduction, under basal conditions or after exposure to LPS+ATP for 3 h. pLex, transduction of lentiviral particles bearing the control vector pLex. **B**. Stable constitutive expression of ORF9b strongly induces caspase-1 activity in A549 lung carcinoma cells. A549 cells stably integrating and constitutively expressing the indicated SARS-CoV-2 accessory proteins were assayed for specific released caspase-1 activity under basal conditions or after exposure to LPS (100 ng/mL) and ATP (5 mM) for 3 h. pLex, A549 integrating the lentiviral vector pLex with no SARS-CoV-2 insert. **C**. A549 cells with stable integration and expression of ORF9b (A549.ORF9b) were transduced with lentiviruses for the expression of the indicated ORFs and released caspase-1 activity determined 48 h after transduction, under basal conditions or after exposure to LPS+ATP for 3h. pLex, transduction of lentiviral particles bearing the control vector pLex. **D**. The induction of caspase-1 activity by ORF9b is dominant among all accessory proteins when concomitantly expressed in A549wt or A549.ORF9b cells. The concomitant expression of all accessory proteins abrogates the LPS+ATP-induced additive induction of caspase-1 activity to that induced by ORF9b alone. Parental A549wt or A549.ORF9b cells were transduced with equal titers of lentiviral particles for each accessory protein and assayed for released caspase-1 activity after 48 h. pLex, transduction of lentiviral particles bearing the control vector pLex. **E**. The induction of caspase-1 activity by the concomitant expression of all SARS-CoV-2 accessory proteins in A549wt cells, without or with additional boosting by LPS+ATP, is dependent on the NLRP3 inflammasome. Parental A549wt cells were transduced with equal titers of lentiviral particles for each accessory protein and assayed for released caspase-1 activity after 48 h. Cells were treated with the NLRP3 inflammasome inhibitor MCC950 for the last 3 h or vehicle (control).

Transient transfection of THP-1 monocytic cells with ORF9b, but not any other SARS-CoV-2 accessory protein except for a more modest induction by ORF7a, also strongly induced caspase-1 activity in the absence of priming signals (Fig. 1C) in a response that was additive to that induced by LPS+ATP (Fig. 1C). In good concordance with the response observed in A549 cells, transient transduction of ORF3b or ORF7a into THP-1 cells inhibited basal caspase-1 activity, and transduction of ORF3a, ORF7a or ORF10 completely abrogated the induction of caspase-1 by LPS+ATP (Fig. 1C).

To examine if the observed inhibition of basal and LPS+ATP-induced caspase-1 activity by ORF3a, ORF7a, ORF8 and ORF10 affected the activation of caspase-1 by ORF9b, we transiently transduced A549 cells stably integrating ORF9b (A549.ORF9b), which displays high basal caspase-1 activity relative to A549.pLex cells, with lentiviral particles for the expression of each of the other accessory proteins or their control vector, pLex. We observed that ORF3a and ORF6, but not ORF7a, ORF8 or ORF10, significantly attenuated the caspase-1 activity induced by ORF9b (Fig. 1D). Interestingly, expression of ORF10 enhanced the caspase-1 activity induced by ORF9b in A549 cells (Fig. 1D). Remarkably, transient transduction of any of the other accessory proteins blunted the caspase-1 activity induced by LPS+ATP in A549.ORF9b cells (Fig. 1D), to a greater extent particularly by ORF3a and ORF7b, but also by ORF8 and ORF10 (Fig. 1D). Transient transduction of THP-1.ORF9b cells, with stable integration and constitutive expression of ORF9B, with any of the other SARS-CoV-2 accessory proteins did not significantly impact basal caspase-1 activity (Fig. 1E). However, similar to the responses seen in A549.ORF9b cells, transduction of any of the other accessory proteins strongly inhibited the caspase-1 activity induced by LPS+ATP in THP-1.ORF9b cells (Fig. 1E).

As all accessory proteins are concomitantly expressed upon infection by SARS-CoV-2, we simultaneously expressed all the accessory proteins under study in parental A549 cells (A549wt), treated, or not, with LPS+ATP after transient transduction of ORFs (48 h). In parallel, we also subjected stable A549.OR9b cells to transient co-transductions with lentiviruses for the expression of the rest of accessory proteins. It should be noted that, in these experiments, each of the 10 accessory proteins is expressed at roughly equimolar levels, i.e., about 1/10 of the expression levels per cell relative to experiments with expression of individual accessory proteins. Concomitant expression of all accessory proteins in A549wt cells strongly (≈ 10-fold) induced caspase-1 activity, while it did not significantly affect the high levels of caspase-1 activity of A549.ORF9b cells (Fig. 1F). Likewise, transient expression of all accessory proteins in A549wt cells, including ORF9b (lane “all ORFs”), did not hinder the induction of caspase-1 activity by LPS+ATP (Fig. 1F), although the induction was attenuated as compared to the expression of ORF9b alone (compare to Fig. 1B, LPS+ATP, lanes “pLex” and “ORF9b”). Moreover, transient expression of all ORFs (except ORF9b) in stable A549.ORF9b cells abrogated the LPS+ATP-induced additive caspase-1 activation (Fig. 1F). These observations suggest that, while some SARS-CoV-2 accessory proteins, notably ORF3a, ORF7b and ORF10, counter inflammasome activation pathways triggered by LPS+ATP, they do not overcome the induction of caspase-1 activity by ORF9b, when expressed at levels likely paralleling those expected from SARS-CoV-2 infection, in which all accessory proteins are presumably expressed simultaneously. In turn, these results suggest that ORF9b and LPS+ATP activate caspase-1 through distinct pathways.

The induction of caspase-1 activation by SARS-CoV-2 accessory proteins was abolished by the NLRP3 inflammasome inhibitor, MCC950, both without and with LPS+ATP (Fig. 1G), indicating that SARS-CoV-2 ORF9b induces caspase-1 activation through the NLRP3 inflammasome. Furthermore, these results suggest that the signals from ORF9b and LPS+ATP likely act divergently upstream of the NLRP3 inflammasome.

### Virtual screening to identify ORF9b dimerization inhibitor compounds

The strong induction of inflammasome-dependent caspase-1 activity by ORF9b led us to hypothesize that this accessory protein might be a major driver of the unbridled cytokine release and inflammation associated with severe COVID-19 ^20^. This prompted us to develop small molecule inhibitors targeting ORF9b, potentially capable of mitigating the excessive activation of the inflammasome and consequent cytokine storm induced by SARS-CoV-2.

ORF9b can occur as monomers or homodimers ^27^ and, while monomers interact with TOM70 ^5,25–29^, in competition with HSP90 ^26,27,30^, ORF9b homodimers have been reported to directly associate with membranes ^29,33^. After modeling of a solved structure of the ORF9b homodimer (PDB:6Z4U), we conducted a blind docking screening on the entire homodimerization surfaces of the proteins, using as ligand sources 96,096 compounds from the European Chemical Biology Diversity Library and 1,056 fragments from the ECBL Fragment-Based lead discovery library. The dimeric ORF9b structure is constructed with 29 hydrogen bonds between two protomers and two salt bridges formed by residue R58 of one protomer and residue E90 of the other protomer ^29^. Following an iterative process of molecular dynamics, docking and energy scoring, a total of 10 ligands were selected (Fig. 2) based on their free binding affinity energies on the ORF9b protein (Table S1), position of the ligand binding sites on the homodimerization surface of the homodimer complex, and commercial availability. The selected ligands were also predicted to interact with favorable energies with the ORF9b monomer, on a surface involved both in homodimerization and in binding to TOM70 ^26^. As such, the use by ORF9b of the same surface for the formation of homodimers and also for interaction with TOM70 implies that these are mutually exclusive interactions ^26^ and, as a corollary, that compounds binding to this surface may potentially prevent both the formation of ORF9b homodimers and its interaction with TOM70.

**Figure 2.**
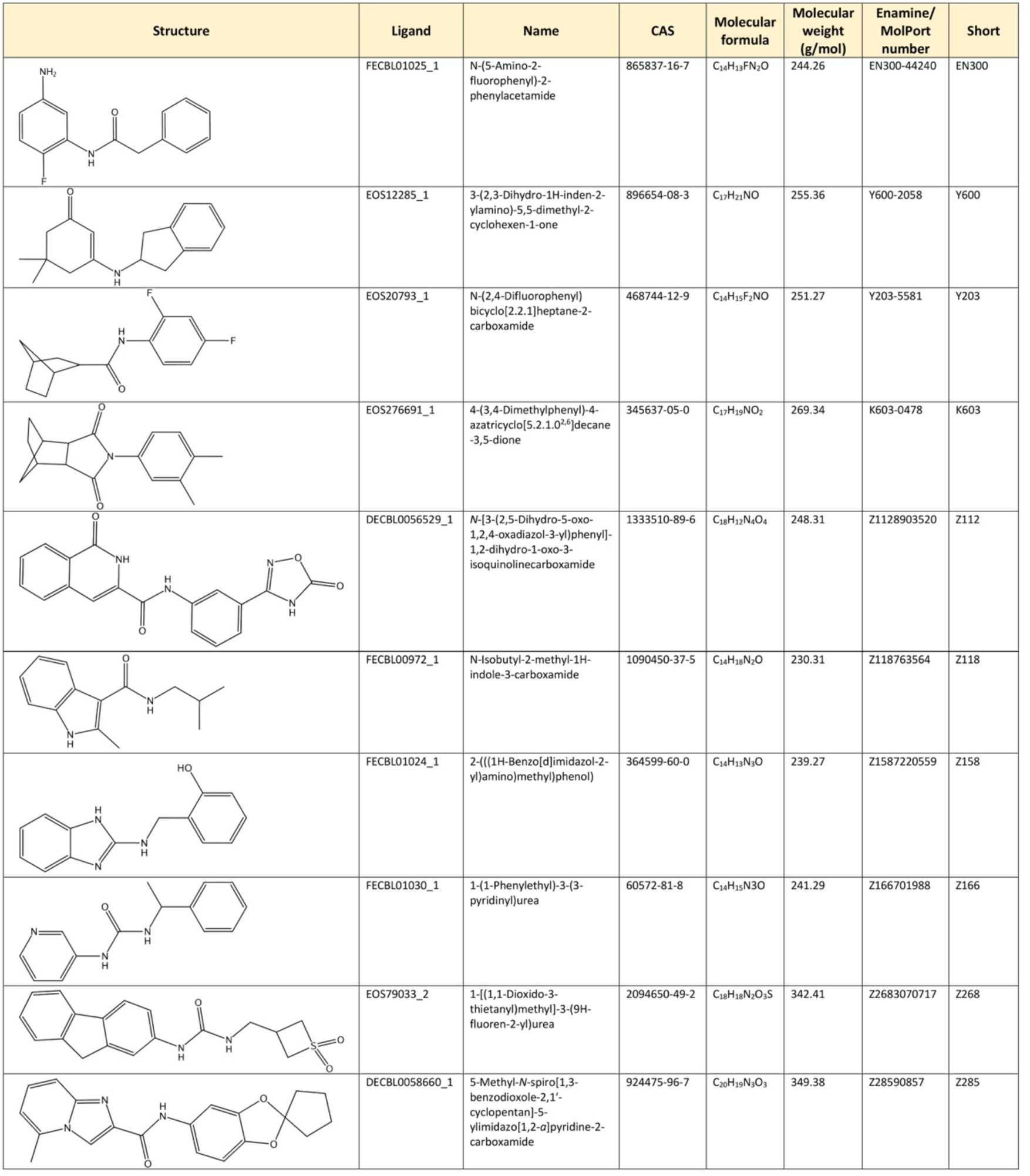
Structures and nomenclature of hit Compound interactors on the ORF9b homodimerization surface. **Column 1**, 2D structure to identify the different scaffolds appearing in the selected compounds. **Column 2**, reference name to locate the compounds in the different virtual databases. The number after the low bar indicates its position in the docking process. **Column 3**, IUPAC name. **Column 4**, CAS Registry Number. **Column 5 and 6**, molecular formula and molecular weight. **Column 7**, compound ID. **Column 8**, compound short name used in the text.

### ORF9b dimerization inhibitors abrogate caspase-1 activation and cytokine release induced by ORF9b

We next experimentally tested the ability of 9 of the 11 compounds selected by virtual screening as potential ORF9b ligands, to prevent homotypic ORF9b interactions, as determined by surface plasmon resonance. For this, recombinant GST-ORF9b was immobilized on CM5 chips and free GST-ORF9b protein, with or without pre-incubation with 5 μM of each of the compounds, allowed to interact in the mobile phase with the GST-ORF9b-derivatized chips. As a control, GST was used in the mobile phase in place of chimeric GST-ORF9b. Binding of GST-ORF9b to immobilized GST-ORF9b was significantly inhibited by two of the Compounds, EN300 and Y600 (Fig. 3A). These two Compounds were among the highest ranking in terms of favorable binding energies to ORF9b in the virtual screening (Fig. 3B, Table S1).

**Figure 3.**
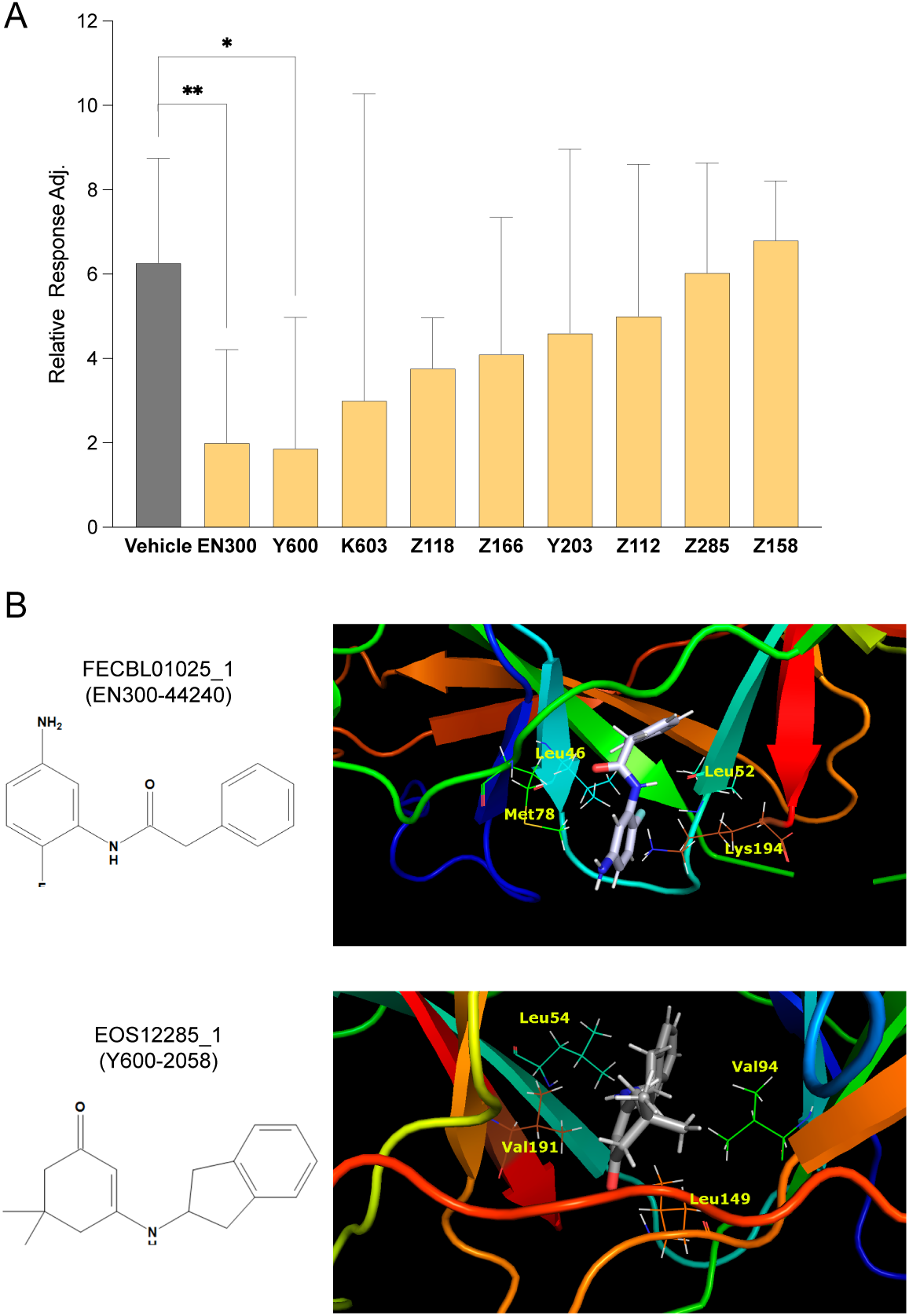
Small Compounds are effective inhibitors of ORF9b-ORF9b homotypic interaction. **A,** Surface plasmon resonance (SPR) determinations of ORF9b homodimerization surface-binding hit Compounds on ORF9b-ORF9b interactions. Recombinant GST-ORF9b protein was immobilized on CM5 chips and interactions with mobile-phase GST-ORF9b Compounds recorded by SPR, in the absence (vehicle) or presence of the indicated hit Compounds at 5 μM final concentration. Determinations were performed in triplicate. **B,** Structures and ribbon models of the two most experimentally active Compounds in inhibition of ORF9b homodimerization interaction, FECBL01015_1 (EN300) and EOS1228_1 (Y600). For the ribbon rendering, the most relevant residues for on ORF9b for its interaction with each Compound are illustrated, all located on the homodimerization surface.

We have shown above that ORF9b is a strong inducer of caspase-1 activation. As such, we tested the ability of EN300 and Y600 to affect this function. Both Compounds, but not Y203, which does not significantly prevent the homotypic binding of recombinant ORF9b, inhibited the activation of caspase-1 in A549.ORF9b and THP-1.ORF9b cells at concentrations in the nanomolar range (Fig. 4A,B). Moreover, EN300 and Y600 caused a rapid and remarkable eviction of ORF9b from mitochondria and relocalization to cytoskeletal structures (Fig. 4C). Collectively, these observations indicate that EN300 and Y600 inhibit the dimerization of ORF9b, thereby preventing its mitochondrial localization, activation of inflammasome-mediated caspase-1 and secretion of pro-inflammatory cytokines.

**Figure 4.**
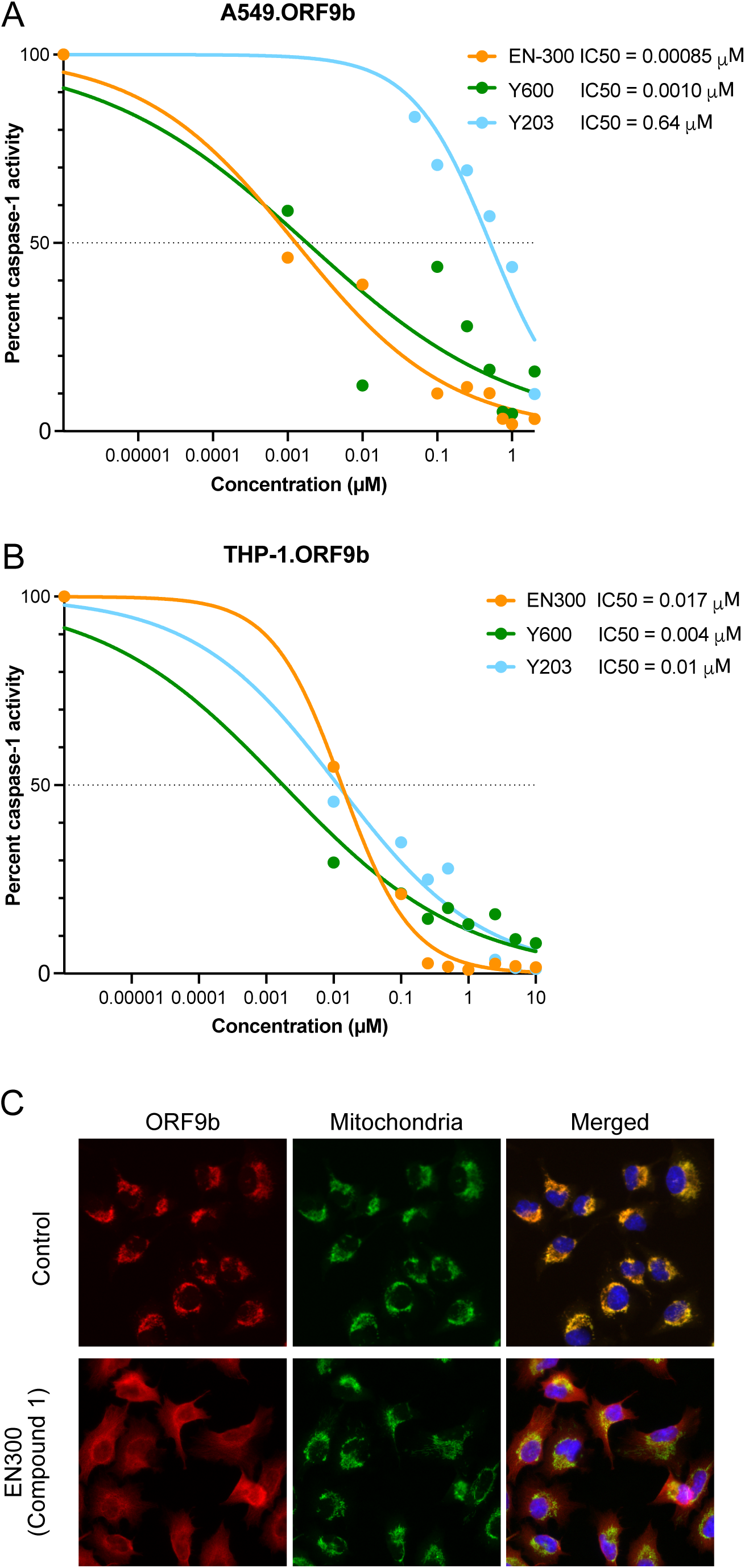
ORF9b homodimerization inhibition Compounds inhibit ORF9b-mediated activities. **A**, Effect of Compounds EN300, Y600 and Y203 on of caspase-1 activity induced by ORF9b in A549 lung epithelial cells. **B**, Effect of Compounds EN300, Y600 and Y203 on of caspase-1 activity induced by ORF9b in THP-1 lung epithelial cells. **C**, Compound EN300 induces mitochondrial eviction of ORF9b in A549.ORF9b cells.

### EN300-similar ORF9b inhibitors

Given the strong activity of compound EN300 on ORF9b homotypic interaction, ORF9b-induced caspase-1 activation, cytokine release and mitochondrial localization, we explored the chemical space to identify further ligands with similar structural features that might provide us with structural diversity which may aid in subsequent chemical optimization. To this end, we conducted a round of constrained docking specifically overlapping with EN300 in the EN300-ORF9b complex model, using the SciFinder database as a source of ligands. Iterations of selection on the basis of free binding energies, classical molecular dynamics simulations and binding energy rescoring, yielded five EN300-similar ligands, designated Compounds 2 through 8 (Fig. 5, Table S2). Hereafter, compound EN300 is designated Compound 1.

**Figure 5.**
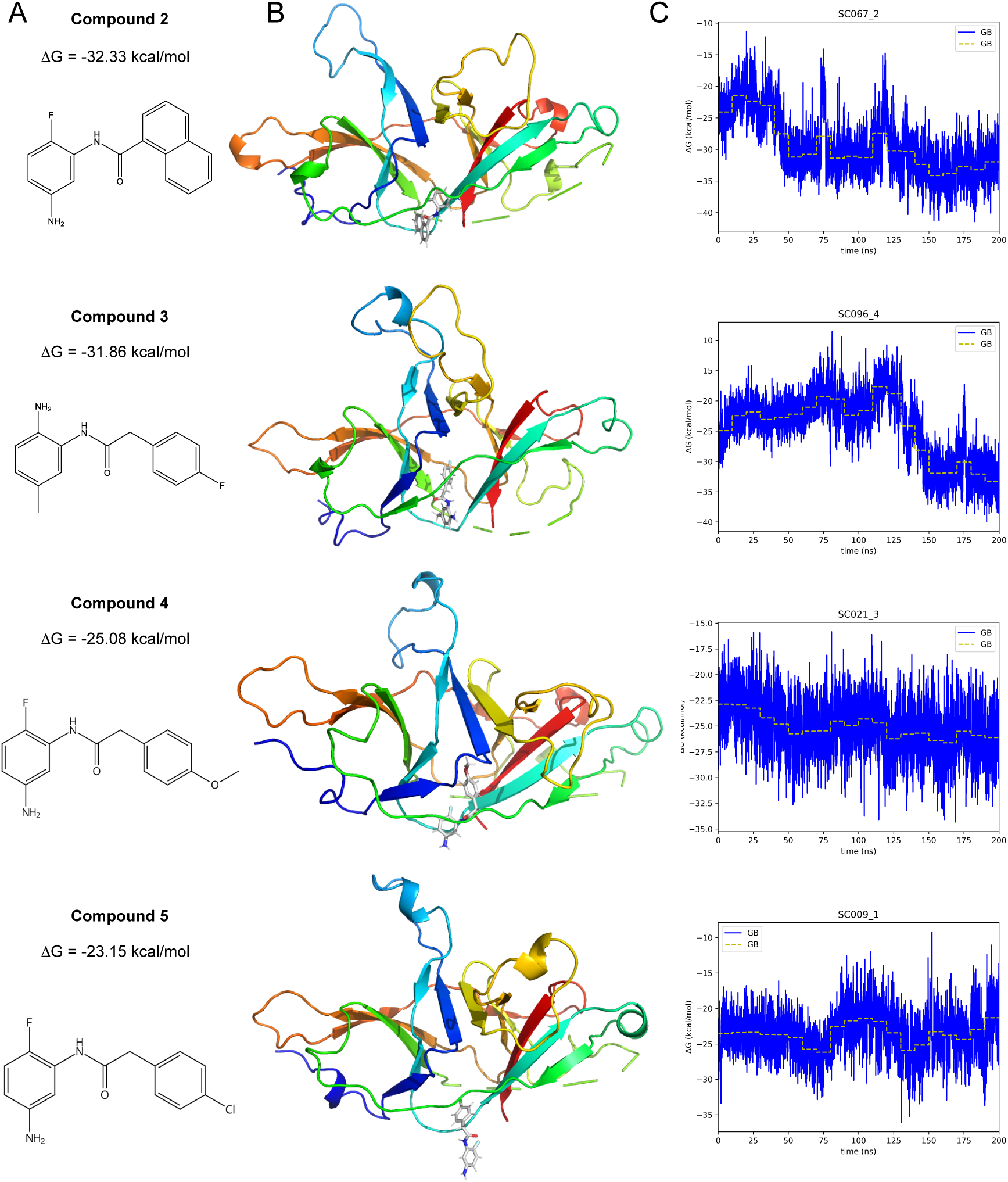
**A,** Structures, ribbon models and molecular dynamics determinations for four EN300-similar compounds, in interaction with the ORF9b homodimer. **A**. 2D structure of the four purchased compounds similar to the EN300 initial hit. Predicted binding free energy using MMGBSA approach, as an average over the last 10 ns of molecular dynamics. **B**. A 3D representation of the predicted binding mode of the compounds with the ORF9B homodimer. **C**. Evolution of the binding free energy, obtained with the MMGBSA methodology, along the time during the molecular dynamics.

These structures served as guides for *de novo* synthesis of the corresponding compounds, as well as additional compounds with small modifications and synthetic intermediate compounds (designated Compounds 6-8). These seven new compounds were analyzed for their ability to inhibit ORF9b-induced activation of caspase-1 and cytokine release, and their ability to cause mitochondrial eviction of ORF9b. Compounds 2, 4, 5, 6 and 7 inhibited ORF9b-induced caspase-1 activation in A549 (Fig. 6A) and THP-1 (Fig. 6B) cells and cytokine release in THP-1.ORF9b cells (Fig. 6C), albeit with lower potency as compared to Compound 1. Except for Compound 4, they were also capable of causing mitochondrial eviction of ORF9b in A549.ORF9b cells at low micromolar concentrations (Fig. 6D). Compounds 3 and 8 exhibited much lower or no ORF9b-induced caspase-1 inhibitory activity in A549 or THP-1 cells or mitochondrial eviction of ORF9b in A549.ORF9b cells (Fig. 6).

**Figure 6.**
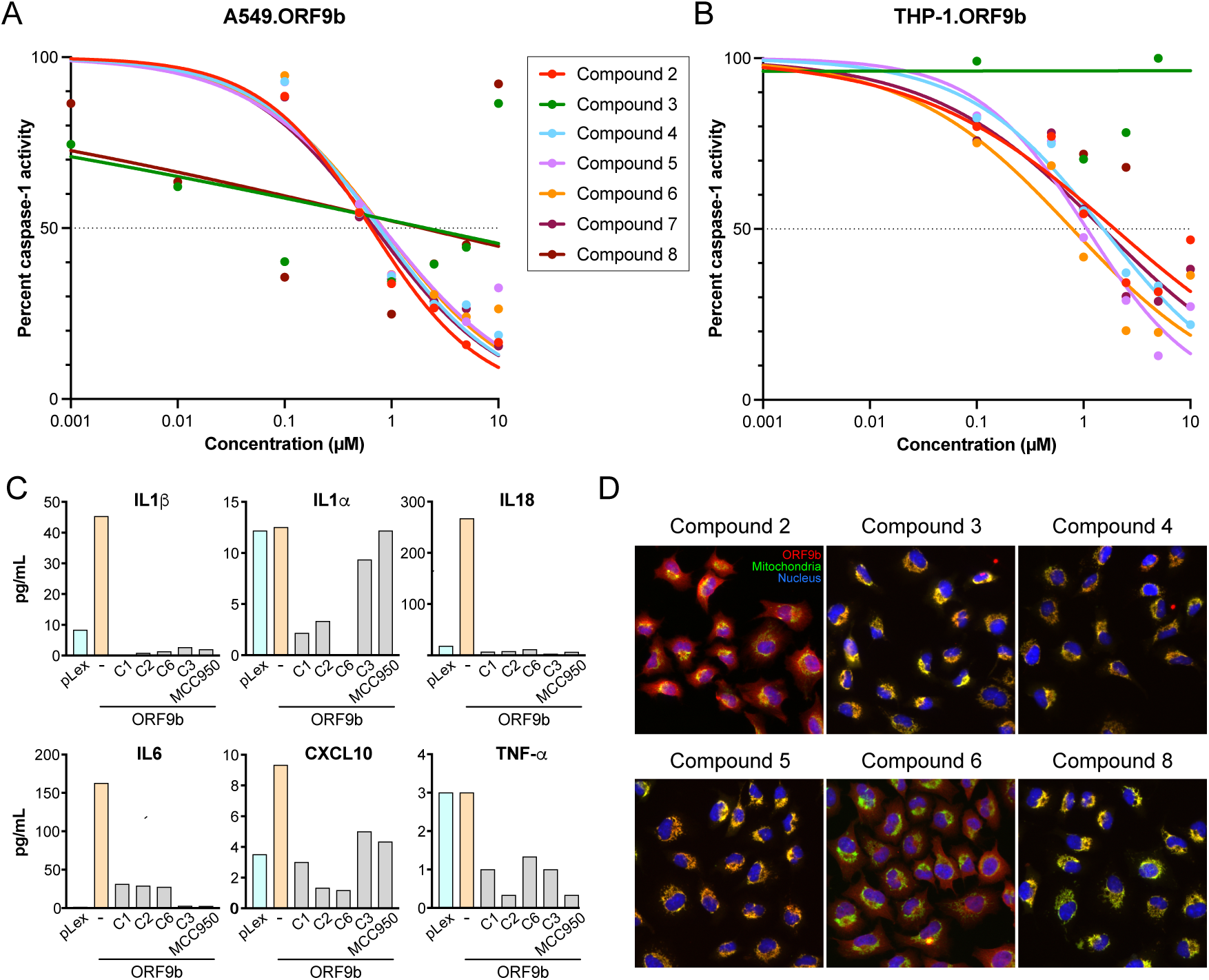
EN300-similar Compounds inhibit ORF9b-mediated activities. **A**, Compounds 2, 4, 5, 6 and 7, but not Compounds 3 and 8, inhibit the activation of caspase-1 induced by ORF9b in A549 lung epithelial cells at low micromolar concentrations. **B**, Compounds 2, 4, 5, 6 and 7, but not Compounds 3 and 8, inhibit the activation of caspase-1 induced by ORF9b in THP-1 monocyte-macrophage cells at low micromolar concentrations. **C**, Compounds EN300, 2 and 6 inhibit the secretion of the indicated cytokines induced by ORF9b in THP-1 monocyte-macrophage cells. Compound 3 also inhibits de secretion of several cytokines induced by ORF9b, except IL-1α. The NLRP3 inflammasome inhibitor, MCC950, inhibits the ORF9-induced secretion of cytokines in THP-1 cells, except IL-1α. **D**, Compounds 2, 4, 6 and 7, but not Compound 3, induce a loss of mitochondrial localization of ORF9b in A549.ORF9b cells.

The most relevant interactions of Compound 1 on ORF9b are residues L52, Q77 (hydrogen bonds), V15, L46, P51, L52, S53, L54, V76, M78 and K97 (hydrophobic interactions). Major interactions of Compound 2, are L52, L54, Q77, M78 (hydrogen bonds), L46, P51, L52, S53, L54, I74, V76, Q77, M78, K97 (hydrophobic). Major interactions of Compound 4 are L52 (hydrogen bonds), V15, L46, L48, P51, L52, S53, L54, V76, M78 (hydrophobic) (Table S3). Several such residues are critical for the homodimerization of ORF9b, notably P51, S53, V93/94 and K97 ^29^. Also of note, L46, L52 and S53 crucially participate in the interaction of ORF9b with TOM70 ^26^. Among these residues, phosphorylation at S53 prevents the interaction of ORF9 with TOM70 ^28,30,35^. As such, these interactions both mechanistically explain the antagonism of ORF9b homodimerization by Compound 1 and its structural analogs and predict that these compounds are also likely to disrupt the interaction of ORF9b monomers with TOM70.

### ORF9b dimerization inhibitors restore IFN-I responses compromised by ORF9b

ORF9b is one of several SARS-CoV-2 proteins that interfere with type I and III interferon signaling through various mechanisms ^28,31,36–39^. We thus tested the effect of SARS-CoV-2 ORF9b expression in A549 lung carcinoma cells or THP-1 monocyte-macrophage cells and the impact of our ORF9b dimerization inhibitors on (1) interferon-sensitive response element (ISRE)-mediated transcriptional responses to the TLR7/8 agonist, Resiquimod ^40^, or the TLR3 agonist, poly(I:C) ^41^, and (2) transcriptional responses induced by exposure of cells to IFN-β1. Treatment of THP-1 cells with Requisimod induced the expression of the IRES-regulated genes, IFNB1, IFIT1 and CCL5 (Fig. S1). Contrary to expectations, expression of ORF9b did not impair this induction (Fig. S1). Similarly, transfection of THP-1 cells with poly(I:C) strongly induced the expression of IFNB1, IFIT1 and CCL5, which was also not countered by ORF9b (Fig. S1). On the other hand, exposure of A549.ORF9b or THP-1.ORF9b cells to IFN-β1 strongly stimulated the expression of the IFN-I/III stimulated genes, OAS3 and ISG15, in control cells (Fig. 7A,B). This response was significantly mitigated in cells expressing ORF9b (Fig. 7A,B), and largely restored by the ORF9b inhibitors, Compounds 1, 2 and 6, but not by their structural analog, Compound 3 (Fig. 7C). ISRE-driven transcriptional responses to IFN-β1 follow distinct signaling pathways involving transmembrane cell-surface receptors and JAK1, TYK2 and STAT1/2 activation cascades that activate the IRF9 transcription factor ^42^. The inhibition of this pathway by ORF9b, shown here, has not been described before.

**Figure 7.**
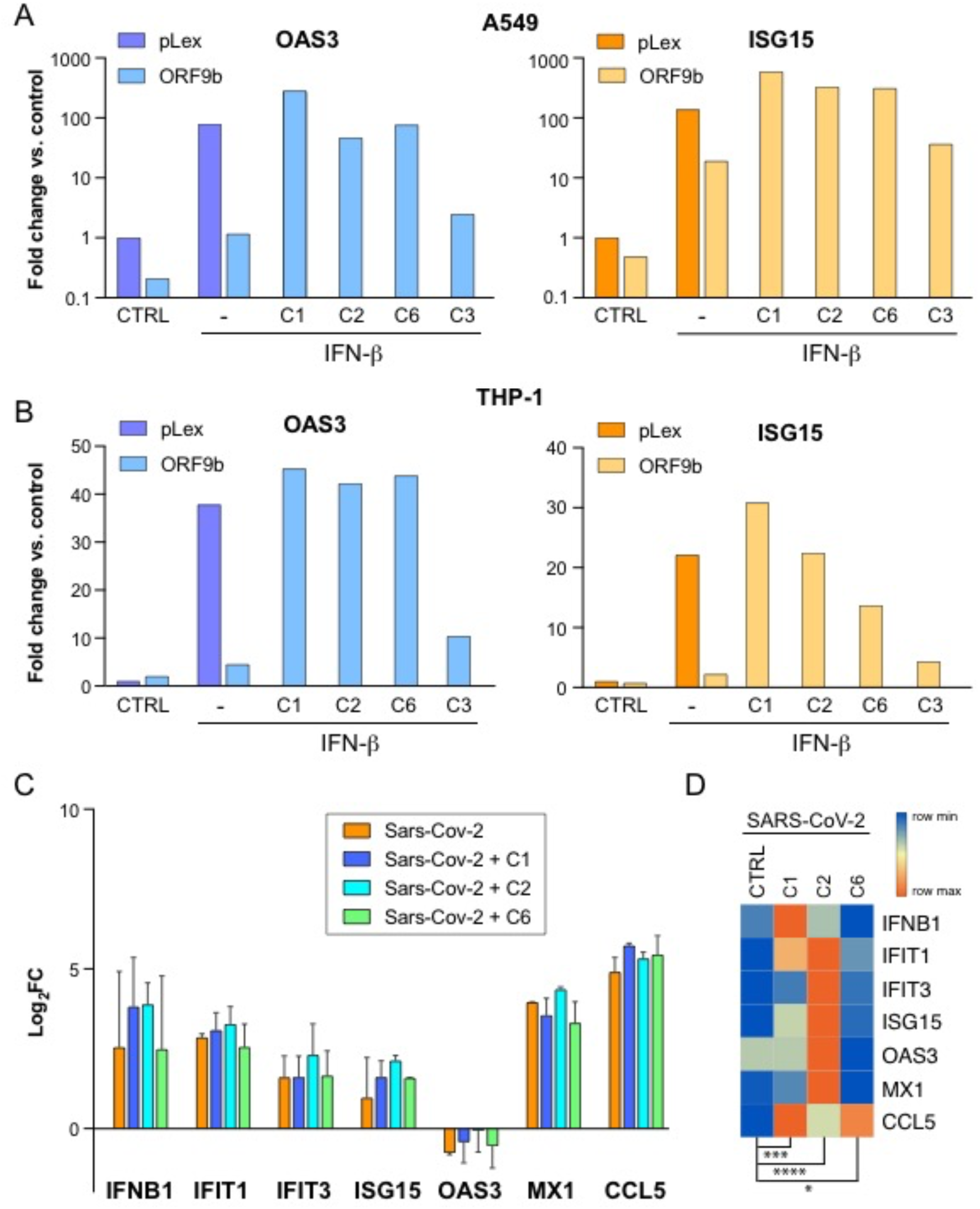
ORF9b dimerization inhibitors restore IFN-I responses compromised by ORF9b. **A**, The ORF9b dimerization inhibitors Compound 1 (C1), Compound 2 (C2) and Compound 6 (C6), but not Compound 3 (C3), restore the IFN-β-induced expression of OAS3 and ISG15 in A549.ORF9b cells. pLex, A549 cells stably transduced with the control vector pLex. The Y axis corresponds to fold-change values of RT-PCR ΔΔCt values for the indicated transcripts. **B**, The ORF9b dimerization inhibitors Compound 1 (C1), Compound 2 (C2) and Compound 6 (C6), but not Compound 3 (C3), restore the IFN-β-induced expression of OAS3 and ISG15 in THP-1 cells transiently transduced with lentiviral particles for the expression of ORF9b or with control lentiviral particles (pLex). The Y axis corresponds to fold-change values of RT-PCR ΔΔCt values for the indicated transcripts. **C**, The ORF9b dimerization inhibitors Compound 1 (C1) and Compound 2 (C2), but not Compound 6 (C6), upregulate IFN-I/III response genes in A549.hACE cells upon infection with SARS-CoV-2. Left panel, histogram of Log_2_FC of RT-PCR ΔΔCt values for the indicated transcripts, in triplicate samples. Right panel, row-normalized heatmap of average RT-PCR ΔΔCt values for the indicated transcripts.

These results indicate that only one of the three IFN-I activation pathways assessed evidenced blunting of responses by ORF9b and subsequent restoration by ORF9b dimerization inhibitors. We next tested the potential impact of these Compounds on IFN-I responses in cells infected with SARS-CoV-2, by infecting A549.hACE2 cells with SARS-CoV-2, with or without incubation with ORF9b dimerization inhibitors. Viral infection moderately induced the expression of IFNB1, IFIT1 and CCL5, as well as ISG15, but not OAS3, which was rather downregulated (Fig. 7C). Treatment of infected cells with the ORF9b dimerization inhibitors, Compounds 1, 2 or 6, using a broad range of concentrations (0.4 to 50 μM) had no impact on the release of viral particles into the culture medium. As a control, the antiviral Remdesivir abrogated viral replication (data not shown). However, and remarkably, Compound 2, but not Compound 6, potentiated the expression levels of IFIT1, IFIT3, OAS3, MX1 and ISG15 in infected cells (Fig. 7C), the differences reaching statistical significance when these genes were considered as a set (Fig. 7D). While Compounds 1 and 2 had no effect on the transcript levels of these genes in uninfected A549.hACE2 cells, Compound 6 counterintuitively induced their upregulation in the absence of SARS-CoV-2 infection (Fig. S1).

## Discussion

A major factor underlying the high pathogenicity of several viruses is an unbridled inflammatory response that can be triggered subsequent to viral infection, in the form of an excessive activation of pro-inflammatory functions of immune cells, including the release of high levels of pro-inflammatory cytokines that may cause multisystemic organ damage unless timely countered by anti-inflammatory interventions ^20^. Such imbalanced responses are not necessarily correlated with the efficiency of viral replication within infected cells ^43^ and are rather driven by viral factors that dysregulate cellular innate immune responses, mainly by interfering with type-I/III interferon (IFN-I/III) pathways and by activating the inflammasome ^44^. While the evolutionary advantage to viral propagation of blunting IFN-I/III antiviral responses by virus-coded proteins is well understood ^45^, that of unabated inflammasome activation is less evident.

SARS-CoV-2 evolved from the ancestral strain that originated in Wuhan, China, to enhance human-to-human propagation and to escape human innate and adaptive immune responses. Viral propagation is optimized mainly by virtue of mutations in the SARS-CoV-2 spike protein that enhance viral binding to its human receptor, angiotensin-converting enzyme 2 (ACE2). However, additional non-synonymous mutations in non-structural, structural, and accessory proteins also affect viral propagation by altering the host response to infection, at least partly through the modulation of ORF9b, ORF6 or N protein levels ^28,43^ and mutations affecting their interactions with cellular proteins that regulate innate immune responses ^43^.

Among SARS-CoV-2 accessory proteins, ORF6 most profoundly antagonizes the antiviral response ^36,37,46–48^, by binding to the nuclear pore complex components RAE1 and NUP98 and inhibiting the nuclear import of STAT-1 and IRF3 ^49–51^, and by inhibiting the nucleocytoplasmic export of mRNAs encoding IFN-I/III pathway factors, including IRF1 and RIG-I ^52,53^, and pro-inflammatory pathway factors, such as NF-κB ^54^. Although less potently than ORF6, SARS-CoV-2 accessory proteins ORF7b, ORF8 and ORF9b have also been reported to antagonize the IFN-I/III and NF-κB pathways ^25,28,31,32,47,48,55–57^ through various mechanisms. For example, ORF9b has been found to interact with the K63 ubiquitin ligase, NEMO, preventing the K63 polyubiquitination of NF-κB upon viral infection, thus antagonizing the production of IFN-β1 and pro-inflammatory cytokines ^32^.

In contrast to the blunting of IFN-I/III pathways, SARS-CoV-2 viral factors trigger a pro-inflammatory program through the activation of inflammasomes ^19,20,22,58^, particularly the NLRP3 inflammasome ^58,59^. The unbridled activation of this pathway may underlie the multi-organ immunopathology caused by the most pathogenic variants of this virus, leading to severe disease and death ^60,61^, a mechanism shared with other highly pathogenic viruses ^62^. SARS-CoV-2 structural (S, N, E), non-structural (nsp1, nsp6, nsp13) and accessory (ORF3a, ORF8) proteins have been shown to affect the activation state of the NLRP3 inflammasome to various degrees and through diverse mechanisms ^63^.

In this study, we have compared nine SARS-CoV-2 accessory proteins, namely ORF3a, ORF3b, ORF6, ORF7a, ORF7b, ORF8, ORF9b, ORF9c and ORF10, for their ability to induce the NLRP3-dependent activation of caspase-1 in the absence of priming signals. We have found that the relative order of intensity of induction of caspase-1 activity by short-term expression (transient transduction) of SARS-CoV-2 accessory proteins is ORF9b >> ORF7b > ORF9c > ORF3a ≈ ORF6 ≈ ORF8 ≈ ORF10. Upon long-term expression (stable integration and constitutive expression), only ORF9b and ORF6 significantly induced caspase-1 activity, in the absence of priming signals. Interestingly, the induction of caspase-1 activity by LPS+ATP was additive to that of ORF9b. Two other accessory proteins, ORF7b and ORF6, also activated caspase-1 additively to LPS+ATP, but only under transient expression. This suggests that these accessory proteins use pathways parallel to those used by LPS+ATP to activate caspase-1. The activation of caspase-1 by ORF9c and ORF3a was not additive to that of LPS+ATP, suggesting that they use overlapping pathways to activate caspase-1.

Remarkably, all other SARS-CoV-2 accessory proteins inhibited the activation of caspase-1 induced by LPS+ATP, particularly ORF7a, which also strongly inhibited basal caspase-1 activity, and also ORF3a, which modestly activated caspase-1 under basal conditions and ORF10, which did not inhibit basal caspase-1 activity. Consistently, ORF3a, ORF7a and ORF10 countered the additional activation of caspase-1 activity induced by LPS+ATP over that induced by ORF9b. We conclude that these three SARS-CoV-2 accessory proteins inhibit the LPS+ATP-mediated activation of caspase-1. However, of these three proteins, only ORF3a, but not ORF7a or ORF10, antagonized the ORF9b-stimulated activation of caspase-1. These results further reinforce the conclusion that ORF9b and LPS+ATP activate NLRP3 inflammasome-dependent caspase-1 through divergent pathways. Although the current study has not explored the specific pathways used by ORF9b to activate caspase-1, its prominent association with, and disruption of, mitochondria ^64–66^ through its interaction with the mitochondrial translocase, TOM70 ^6,26,27^, suggests a mitochondrial pathway of NLRP3 inflammasome activation ^17^ as a prime candidate pathway.

Importantly, we have found that the strong activation of caspase-1 by ORF9b overrides the combined inhibitory activities of all other accessory proteins expressed together, attesting to the potency of inflammasome activation by ORF9b, and highlighting this accessory protein as a relevant therapeutic target to counter the exacerbated pro-inflammatory responses triggered by highly immunopathogenic SARS-CoV-2 variants. To that end, we have conducted a virtual screening for small molecules that has yielded small molecules that inhibit the homodimerization of ORF9b and, as inferred from the eviction of ORF9b from mitochondria by these compounds, also likely inhibiting its interaction with TOM70. The most active of these compounds potently countered the activation of caspase-1 in A549 lung cancer and THP-1 monocyte-macrophage cells and the release of pro-inflammatory cytokines in THP-1 cells induced by ORF9b. Contrary to other reports ^25,31^, we did not observe an antagonism by ORF9b of the activation of the IFN-I pathway by the TLR3 and RIG-I ligand, poly(I:C), or by the TLR7/8 agonist, Resiquimod. However, ORF9b did significantly inhibit the transcriptional response to exposure of A549 and THP-1 cells to exogenous IFN-β1, suggestive of a previously unknown mechanism of antagonism of IFN-I signaling by ORF9b. The same ORF9b homodimerization inhibitors that countered the activation of caspase-1 by ORF9b, namely Compounds 1, 2 and 6, restored the transcriptional response to IFN-β1 abrogated by ORF9b in A549 and THP-1 cells. Importantly, these compounds, particularly Compound 2, enhanced the IFN-I response to infection of A549.hACE2 cells with SARS-CoV-2. Of interest, SARS-CoV-2 infection caused the upregulation of the IFN-I pathway target genes, IFNB1 and ISG15, and a downregulation of OAS3. Compound 2 reverted this downregulation.

In conclusion, we have discovered novel small molecule inhibitors of ORF9b homodimerization that mitigate the excessive inflammasome activation and the abrogation of IFN-I signaling mediated by ORF9b. These compounds could be useful additions to the currently available therapeutic arsenal against COVID-19, by boosting antiviral responses while, simultaneously, moderating excessive pro-inflammatory responses conducive to the immunopathology and disease severity provoked by SARS-CoV-2 in many patients. A limitation of this study is that, while robust evidence was produced in cell lines, we have not demonstrated the inflammatory mitigating activities of the new compounds upon SARS-CoV-2 infection *in vivo*. Nevertheless, the results presented here warrant the future completion of all pertinent pre-clinical demonstrations, including animal models of COVID-19.

## Materials and Methods

### Lentiviral transduction

A549 pulmonary epithelial cells and THP-1 cells were cultured in Dulbecco’s Modified Eagle Medium (DMEM) (Gibco) supplemented with 10% (v/v) heat-inactivated fetal bovine serum (FBS) (Gibco), 1% Penicillin-Streptomycin (100 U/ml) (Gibco) and Amphotericin B (Gibco). SARS-CoV-2 (Wuhan-Hu-1 isolate) accessory proteins coding sequences (codon-optimized for mammalian expression) were cloned into pLVX-EF1α-IRES-Puro Cloning and Expression Lentivector (Clontech) to generate pseudotyped lentiviral particles ^66,67^. Accessory proteins were C-terminally 2 x Strep-tagged to check viral protein expression. For transient experiments, transduction was performed at a multiplicity of infection (MoI) of ≈ 10 to ensure expression of the transduced viral proteins in all cells, followed by experimental perturbations 48 h after transduction. For stable Integration of SARS-CoV-2 accessory proteins, A549 cells were exposed to puromycin (2 µg/mL) 24 h after transduction and selected for 3 d. All cells were cultured at 37 °C in a 5% CO_2_, 90% humidity atmosphere.

### Caspase-1 activity

Released caspase-1 enzymatic activity of was quantitated with the Caspase-Glo 1 inflammasome assay (Promega), following instructions from the supplier. Briefly, cells were seeded, subjected to experimental perturbations, and 50 μL of cell culture supernatant was added to a mixture containing the selective caspase-1 substrate, Z-WEHD-aminoluciferin, and MG-132 protease inhibitor, with or without the irreversible caspase-1 inhibitor, Z-YVAD-FMK (Sigma-Aldrich), and luminescence measured on a Cytation 5 instrument (Agilent). The luminescence inhibited by Z-YVAD-FMK corresponds to specific caspase-1 activity. All assays were done in triplicate.

### Cytokine quantification

Cytokines (IL-1α, IL-1RA, IL-1β, IL-18, IL-6, CXCL10, TNF-α) were quantified by means of Procartaplex multiplex assays (InVitrogen, ThermoFisher), following instructions by the supplier. Briefly, capture beads were incubated for 2 h with cleared cell supernatants, washed and incubated with biotinylated detection antibody mix, incubated for 30 min, washed and incubated with Streptavidin-PE for 30 min, washed and data acquired on a Luminex 200 instrument.

### Virtual screening for small molecule ORF9b homodimerization inhibitors

The atomic coordinates of the solved structure of the SARS-CoV-2 ORF9b homodimer (PDB: 6Z4U) were used as the initial scaffold for virtual screening. This structure contains 194 amino acids, and its X-ray structure is missing several residues of the protein. To complete the system, Modeller ^68^ was used to model the missing residues without significantly affecting the structure of the entire the protein. Subsequently, pH was adjusted to 7.4 via protonation. A monomeric structure with a total of 97 amino acids was retrieved from the dimeric structure. Both structures underwent a 4-step minimization process, as follows: (1) The entire system was fixated except for the added amino acids, allowing the focus to be on the regions of interest without affecting the rest of the structure. (2) The protein backbone was fixated, allowing adjustments of the side chains but not the main structure, which remained unchanged. (3) Reduction of the applied force constant to allow greater flexibility in the atoms during optimization. (4) Removal of all restraints to relax the system, seeking its minimum free energy. Subsequently, The force fields ff14SB and gaff2 were used to describe the protein and the ligands respectively as included in the AMBER20 software. The TIP3P water force field was used for solvation.

Multiple classical and accelerated molecular dynamics (GaMD) simulations ^69^ were performed at different initial temperatures. This temperature change ensures that each dynamics starts with a different velocity distribution, which can have a significant impact on the behavior and properties during the simulation and, thus, explore different states and trajectories of the system. The first step consisted in slowly heating the system under NTV conditions; and perform small constant pressure simulations to gradually adapt the system density. Once the system was equilibrated, a 1,000 ns production simulation was performed, followed by clustering. Both Root Mean Square Deviation (RMSD) and Root Mean Square Fluctuation (RMSF) were calculated for all dynamics and referenced to the same structure. An iterative process of 3 repetitions was performed to guarantee variability in the clusters and their respective representatives. To analyze the conformational space explored by the dynamics, said dynamics were represented along with the cluster representatives in the covariance matrix projections. Representatives with greater diversity and an occupancy greater than 10% were selected.

A direct docking process was conducted with the AutoDock Vina software ^70,71^, using as ligand sources the Diversity Library - European Chemical Biology Library (https://www.eu-openscreen.eu/services/Compound-collection/european-chemical-biology-library-ecbl-diversity-library.html), with 96096 diverse Compounds, the Fragment Library - Fragment-Based lead discovery, with 1056 fragments from the ECBL library (https://www.eu-openscreen.eu/services/Compound-collection/fragment-library-fbld.html) and the SciFinder database (https://www.cas.org). Ligand poses were selected based on energy thresholds, the systems minimized, and the free binding energy calculated using the MMPBSA method. Structures present in the largest number of representative proteins were chosen, and molecular dynamics simulations performed. The systems were selected through an iterative process of molecular dynamics production, energy rescoring and ligand selection. A total of 500 ns with cMD and 400 ns with GaMD were simulated. This process was replicated for both the dimer and the monomer.

### Surface plasmon resonance

Recombinant SARS-CoV-2 GST-ORF9b protein (BPS Bioscience) was immobilized on a Sensor Chip CM5 channel (Cytiva). Control GST protein was immobilized on a second channel of the same chip. For the mobile phase, GST-ORF9b was maintained at 400 nM in running buffer (10 mM HEPES, 0.15 M NaCl, pH 7.4). To assess its activity on the homotypic interaction with solid-phase GST-ORF9b, mobile-phase GST-ORF9b was preincubated with the test Compounds at varying concentrations, and interaction assays run on a Biacore T100 instrument.

### Fluorescence microscopy

Cultured cells were seeded on 96-well flat bottom microplates (Greiner 655090) and allowed to attach for 24 h. Cells were washed with PBS 1x, fixed with 4% paraformaldehyde/PBS during 15 min and, after washing with PBS 1x, the fixed cells were blocked with 1% BSA/PBS and 0.1% Triton X-100, followed by incubation with primary antibodies (rabbit polyclonal anti-ORF9b, Abyntek #9191 and rabbit polyclonal anti-TOMM70, Invitrogen #PA5-83890) diluted in blocking buffer during 1 h. Subsequently, after washing with PBS 1x, cells were incubated with secondary antibody (AlexaFluor-568-conjugated donkey anti-rabbit IgG(H+L), Invitrogen) 1:2000 in blocking buffer containing also 4,6-diamidino-2-phenylindole (DAPI, 1/10,000 dilution), for 1 h at room temperature on a shaker. For mitochondrial visualization, following fixation, cells were incubated with 100 nM MitoView Green (Biotium) for 1 hour at RT on a shaker, then rinsed with PBS 1x before viewing. All images were captured with a Leica Stellaris 8 Confocal microscope.

### Real-time quantitative RT-PCR

Cells were rinsed with PBS and plates frozen. RNA was isolated using Trizol reagent (Invitrogen). Reverse transcription was carried out with 1 μg RNA at 37 °C for 1 h with the following reagents: 5x reverse transcription buffer (Invitrogen), 0.1 M dithiothreitol (DTT) (Invitrogen), Random Hexamers (Roche), 40 U/μL RNAsin (Promega, Fitchburg, WI, USA), 40 mM dNTPs (Bioline, London, UK), 200 U/μL M-MLV-RT (Invitrogen). Gene expression analysis was performed on an Applied Biosystems Quantstudio 3 Real-Time PCR System, using Taqman gene specific sequences for OAS3 (Hs00196324_m1) and ISG15 (Hs01921425_s1) (Applied Biosystems, Thermo Fisher Scientific Inc.). Reactions were performed in a final volume of 20 μL, containing 9 μL of cDNA mixture and 11 μL of the specific Taqman in Master Mix (Applied Biosystems). Real-Time PCR was conducted according to the following parameters: an initial incubation at 50 °C for 2 min and denaturalization at 95 °C for 10 min, followed by 40 cycles at 95 °C and 60 °C for 15 sec and 1 min, respectively. Expression was quantified by ΔΔCt method using Rna18s5 (Hs03928985_g1, Applied Biosystems) as a reference transcript.

## Additional methods are described in Supporting Information

### Funding Statement

This work was funded by the Spanish National Research Council (CSIC, project numbers CSIC-COV19-006, CSIC-COV-19-201, CSIC-COV-19-117, SGL2103019, SGL2103015, 202020E079 and 202320E187), the Catalan Agency for Management of University and Research Grants (AGAUR, 2020PANDE00048, 2021SGR1490, 2021SGR00350), the Spanish Ministry of Science (PID2021-123399OB-I00), the CSIC’s Global Health Platform (PTI Salud Global), The Networked Center for Biomedical Research in Liver and Digestive Diseases (CIBER-EHD), the Spanish Structures and Excellence María de Maeztu program (CEX2021-001202-M) and the European Commission-Next Generation EU (Regulation EU 2020/2094).

## Acknowledgments

The authors are grateful to Dr. Urtzi Garaigorta and Dr. Pablo Gastaminza for performing experiments involving SARS-CoV-2, Dr. Enjuanes (CNB-CSIC, Spain) for providing the Vero E6 cell line and Dr. Molenkamp (EUMC, Netherlands) for the SARS-CoV-2 viral isolate which was provided through EVA-GLOBAL Repository (grant agreement 871029). We also thank the CNB Antiviral Screening Platform for evaluation of the antiviral activity and cytotoxicity of compounds in a SARS-CoV-2 cell culture infection model described in this manuscript.

## Conflict of interest statement

Erika Zodda, Mònica Pons, Natàlia de Moya Valenzuela, Cristina Calvo González, Cristina Benítez Rodríguez, Blanca Díes López-Ayllón, Achraf Hibot, Marta Cascante, María Dolors Pujol Dilme, María Montoya González, Jaime Rubio Martínez and Timothy M. Thomson are co-inventors in patent application EP24382161.8.

## Notes

### Competing Interest Statement

Erika Zodda, Monica Pons, Natalia DeMoya-Valenzuela, Cristina Calvo Gonzalez, Cristina Benitez-Rodriguez, Blanca Dies Lopez-Ayllon, Achraf Hibot, Marta Cascante, Maria Dolors Pujol, Maria Montoya, Jaime Rubio-Martinez and Timothy M. Thomson are co-inventors in patent application EP24382161.8.

### Summary of Updates

1. Update of corresponding author affiliations. 2. Correction of several grammatical and citation errors.

