## Supplemental Methods, Tables and Figure for "Induction of the inflammasome by the SARS-CoV-2 accessory protein ORF9b, abrogated by small-molecule ORF9b homodimerization inhibitors"

Contents:

Supplemental Materials and Methods
Supplemental Reference
Supplemental Tables ST1-ST3
Supplemental Figure SF1

### Supplemental Materials and Methods

Compound synthesis

*General methods***.** Melting points were obtained in open capillary tubes on an MFB-595010M Gallenkamp apparatus equipped with a digital thermometer. IR spectra were obtained using a FTIR Perkin-Elmer 1600 Infrared Spectrophotometer. Only noteworthy IR absorptions are listed ($cm^{-1}$). $^{1}H$ and $^{13}C$ NMR spectra were recorded at room temperature on a Brucker-400 (400 and 100.6 MHz respectively) or Varian Gemini-400 (400 and 100.6 MHz) instruments using $CDCl_3$ as solvent with tetramethylsilane as internal standard, $(CD_3)_2CO$ or $(CD_3)_2SO$. Other $^{1}H$, $^{13}C$ NMR spectra and heterocorrelation $^{1}H$-$^{13}C$ (HSQC and HMBC) experiments were recorded on a Varian Gemini-400 (400 MHz and 100.6 MHz) instrument. Chemical shifts (δ scale) are reported in parts per million (ppm) relative to the central peak of the solvent (δ = 7.26 ppm for $CDCl_3$ in $^{1}H$ NMR and δ = 77.16 ppm for $CDCl_3$ in $^{13}C$ NMR). Mass spectra were taken on a Hewlett-Packard 5988-A instrument and high-resolution mass spectra (HRMS) were recorded on a LC/MSD-TOF mass spectrometer (Agilent Technologies). Column chromatography was performed with silica gel (E. Merck, 70-230 mesh). Reactions were monitored by TLC using 0.25 mm F-254 silica gel (E. Merck). Elemental analysis for C, H and N were determined on a Carlo Erba–1106 analyzer. All reagents were of commercially quality or were purified before use. Organic solvents were of analytical grade or were purified by standard procedures. Reactions were carried out under argon.

*Method for nitro-amide synthesis. Method A-1.* To a solution of the appropriate aniline (1 eq) in 10 mL of dichloromethane, trimethylamine (1 eq) was added to a solution of the appropriate aniline (1 eq) in 10 mL dichloromethane. Then the appropriate carboxylic acid (1 eq) and thionyl chloride were added slowly to the reaction. and the reaction mixture stirred at room temperature for 24 hours. Finally, the solvent was evaporated under reduced pressure and the product recrystallized with a mixture of ethyl acetate and hexane.

*Method for nitro-amide synthesis. Method A-2.* The corresponding acid chloride (1.1 eq) was added to a solution of the nitroaniline (1 eq) in dichloromethane (20 mL) previously cooled to 0 °C with an external ice bath. After addition, the reaction mixture was kept under stirring for 6 h. Reaction controls were carried out by TLC. Once the starting product had been exhausted, the dichloromethane phase was washed with a 1N NaOH solution (3 x 20 mL) and the organic phase was dried over $Na_2SO_4$. Next, the organic phase was filtered and the solvent evaporated under reduced pressure. The obtained crude product was purified by column chromatography using hexane and ethyl acetate as eluents or crystallized from mixtures of hexane and ethyl acetate.

*Method for nitro-amide synthesis. Method A-3.* $TiCl_4$ (0.738 g, 0.42 mL, 3.891 mmol) and the 5-methyl-2-nitroaniline (0.197 g, 1.297 mmol) were added to a solution of 4-fluorophenylacetic acid (0.2g, 1.297mmol) in pyridine (10 mL). The tightly sealed screw-capped vial containing the reaction mixture was then heated at 120 °C. After magnetic stirring for about 72 h, TLC analysis (AcOEt/Hex 20:80 v/v) of the reaction mixture showed complete conversion of the carboxylic acid precursor. The reaction mixture was then cooled, neutralized with HCl solution (2N). The green precipitate obtained was filtered, washed several times with distilled water and dried at 50 °C.

*Method for reduction of nitro-amides. Method B.* The corresponding nitro-amide (1 eq) was treated with iron powder (10 eq) in acetic acid (10 mL) at reflux for 24 hours. At the end of the TLC-controlled reaction, the crude mixture was filtered under reduced pressure followed by neutralization with a 5% ammonia solution. Ethyl acetate (3 x 20 mL) was used to extract the crude reaction product, the organic phases combined, dried over anhydrous $Na_2SO_4$ and filtered under reduced obtain the product. The aminoamide obtained was purified by column chromatography or by recrystallization with hexane/ethyl acetate/hexane mixtures.

***N*-(5-amino-2-fluorophenyl)-1-naphthamide (Compound 2)**

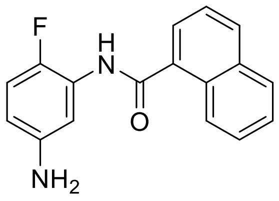


Starting from 200 mg (1.28 mmol) of the 2-fluoro-5-nitroaniline and following the *procedure A-2* was obtained the corresponding amide as a yellow solid in a 67% of yield. Log P = 3.79. NMR $^1$H ($CDCl_3$, 400 MHz) $\delta$ (ppm): 7.30 (t, *J* = 9.2 Hz, 1H, H-7); 7.53-7.60 (m, 2H, H-4, H-6); 7.80 (d, *J* = 7 Hz, 1H, H-3); 7.92 (d, *J* = 7Hz, 1H, H-5); 8.02-8-08 (m, 3H, H-2, H-3', H-6'); 8.40 (d, *J* = 8 Hz, 1H, H-4'); 9.52 (dd, *J* = 2.7, *J* = 9.2 Hz, 1H, H-8). The nitro amide obtained previously 390 mg (1.25 mmol) was reduced following the *procedure B*. After purification by chromatography column (hexane-ethyl acetate 7:3) was obtained the aminoamide **2** in 46% yield. Log P = 3.19. Aspect: brown solid. Mp: 151-153 °C (ethyl acetate) NMR $^1$H ($CDCl_3$, 400 MHz) $\delta$ (ppm): 6.37-6.39 (m, 1H, H-4'); 6.91 (t, *J* = 8.7 Hz, 1H, H-6); 7.50-7.59 (m, 3H, H-3, H-3', H-6'); 7.74 (d, *J* = 7Hz, 1H, H-5); 7.89-7.91 (m, 2H, H-4, H-7); 7.97 (d, *J* = 8 Hz, 1H, H-2); 8.00 (bs, 1H, NH); 8.40 (dd, *J* = 1.0, *J* = 8.0 Hz, 1H, H-8).

***N*-(2-amino-5-methylphenyl)-2-(4-fluorophenyl)acetamide (Compound 3)**

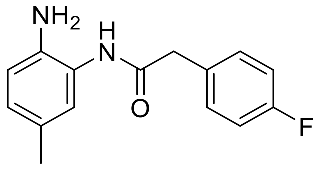


Starting from 130 mg (0.45 mmol) of the corresponding nitroamide **8** and following the general *procedure B* was obtained the desired aminoamide **3** in 36% of yield (43.0 mg). Log P = 2.63. Aspect: brown semi-solid. NMR $^1$H ($CDCl_3$, 400 MHz) δ (ppm): 2.37 (s, 3H, $CH_3$-); 4.08 (s, 2H, $CH_2$-Ar); 6.80 (t, *J* = 6.7 Hz, 2H, H-3, H-5); 7.01 (dd, *J* = 1.5, *J* = 8.2 Hz, 1H, H-4’); 7.09 (dd, *J* = 1, *J* = 6.7 Hz, 2H, H-2, H-6); 7.21 (ba, 1H, H-6’); 7.34 (d, *J* = 8.2 Hz, 1H, H-3’).

**_N_-(5-Amino-2-fluorophenyl)-2-(4-methoxyphenyl)acetamide (Compound 4)**

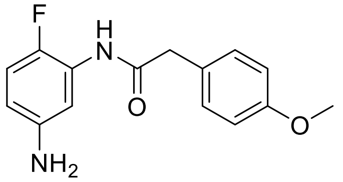


Starting from 100 mg (0.329 mmol) of the nitroamide **7** and following the general *procedure B* was obtained the desired aminoamide **4** in 39% of yield (35.2 mg). Log P = 2.01. Aspect: brown solid. Mp: 147-149 °C NMR $^1$H ($CDCl_3$, 400 MHz) δ (ppm): 3.83 (s, 2H, $CH_2$-Ar); 3.91 (s, 3H, $CH_2$-O); 4.94 (bs, 2H, $NH_2$); 6.95-7.02 (m, 4H, H-3, H-5, H-4’, H-6’); 7.26-7.28 (m, 2H, H-2, H-6); 7.87-7.89 (m, 1H, H-4’); 8.02 (d, *J* = 2.8 Hz, 1H, H-6’); 9.9 (bs, 1H, NH).

**_N_-(5-Amino-2-fluorophenyl)-2-(4-chlorophenyl)acetamide (Compound 5)**

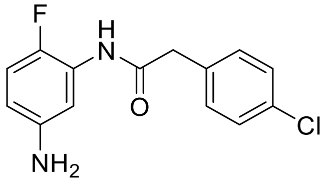


Starting from 100 mg (0.323 mmol) of the corresponding nitroamide **6** and following the general *procedure B* was obtained the desired aminoamide **5** in 42% of yield (37.7 mg). Log P = 2.70. Aspect: brown semi-solid. NMR $^1$H ($CDCl_3$, 400 MHz) δ (ppm): 3.73 (s, 2H, $CH_2$-Ar); 6.98 (t, *J* = 9.2 Hz, 1H, H-3’); 7.27 (d, *J* = 6 Hz, 2H, H-3, H-5); 7.35 (d, *J* = 7 Hz, 2H, H-2, H-6); 7.47 (bs, 2H, $NH_2$); 7.43 (bs, 1H, NH-amide); 7.60-7.63 (m, 1H, H-4’); 8.02 (d, *J* = 2.8 Hz, 1H, H-6’).

**2-(4-Chlorophenyl)-_N_-(2-fluoro-5-nitrophenyl)acetamide (Compound 6)**

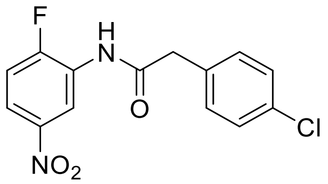

Starting from 183 mg (1.172 mmol) of 2-fluoro-5-nitroaniline and following the general *procedure A-1* was obtained the desired nitroamide **6** in 79% of yield (420 mg). Log P = 3.47. Aspect: white solid. Mp: 224-226 °C. NMR $^1$H ($CDCl_3$, 400 MHz) δ (ppm): 3.78 (s, 2H, $CH_2$-Ar); 7.19 (t, *J* = 9.0 Hz, 1H, H-3'); 7.29 (d, *J* = 7 Hz, 2H, H-3, H-5); 7.40 (d, *J* = 7 Hz, 2H, H-2, H-6); 7.43 (bs, 1H, NH-amide); 7.92-7.96 (m, 1H, H-4'); 9.02 (d, *J* = 2.8 Hz, 1H, H-6').

**_N_-(2-Fluoro-5-nitrophenyl)-2-(4-methoxyphenyl)acetamide (Compound 7)**

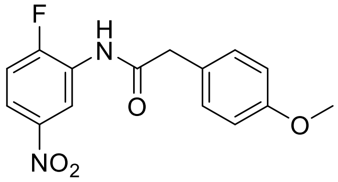

Starting from 470 mg (3.0 mmol) of 2-fluoro-5-nitroaniline and following the general *procedure A-1* was obtained the desired nitroamide **7** in 52% of yield (420 mg). Log P = 2.98. Aspect: white solid. Mp: 181-183 °C. NMR $^1$H ($CDCl_3$, 400 MHz) δ (ppm): 3.76 (s, 2H, $CH_2$-Ar); 3.78 (s, 3H, $CH_3$-O); 6.92 (d, *J* = 6.6 Hz, 2H, H-3, H-5); 7.15 (t, *J* = 9.0 Hz, 1H, H-3'); 7.25 (d, *J* = 7 Hz, 2H, H-2, H-6); 7.60 (bs, 1H, NH-amide); 7.92-7.96 (m, 1H, H-4'); 9.26 (d, *J* = 2.8 Hz, 1H, H-6').

**_N_-(2-Nitro-5-methylphenyl)-2-(4-fluorophenyl)acetamide (Compound 8)**

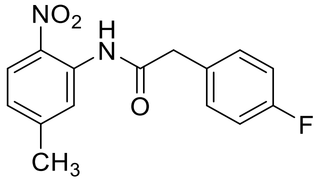

Starting from 197 mg (1.3 mmol) of 2-nitro-5-methylaniline and following the general *procedure A-3* was obtained the desired nitroamide **8** in 98% of yield (367 mg). Log P = 3.27. Aspect: white solid. Mp: 257-259 °C. NMR $^1$H ($CDCl_3$, 400 MHz) δ (ppm): 3.78 (s, 2H, $CH_2$-Ar); 6.01 (bs, 1H, NH); 6.94 (dd, *J* = 1, *J* = 8.6 Hz, 1H, H-4'); 7.11 (t, *J* = 5.2 Hz, 2H, H-3, H-5); 7.13 (dd, *J* = 3.6, *J* = 5.2 Hz, 2H, H-2, H-6); 8.00 (d, *J* = 8.6 Hz, 1H, H-3'); 8.60 (d, *J* = 1 Hz, 1H, H-6').

**Markush formula**

According to the compounds prepared as above, we can establish the following Markush formula:

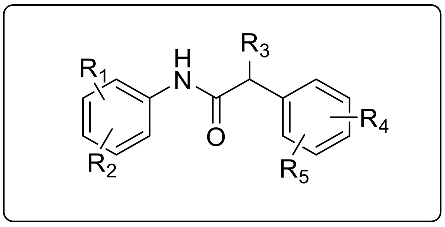


Where the $R_1$ radical is attached to the benzene ring at any of the three possible substitution positions (ortho, meta and para), and via a C atom or heteroatom. Preferred examples of $R_1$ are: -$CH_3$, CN, $CF_3$, CHO, $CH_2OH$, F, H. Other $R_1$ are possible, e.g.: (C1-C3)-alkyl, Cl, Br and I, etc. The $R_2$ radical is attached to the benzene ring at any of the three possible positions (ortho, meta and para), and via a N atom. Preferred examples of $R_2$ are: -$NH_2$, $NO_2$, NHOH, NO, NH-$NH_2$. Other $R_2$ are possible, e.g.: -NH-$COCH_3$, NH-COH, NH-CO-OEt, NH-CO-OMe, NH-CO-$NH_2$ etc. The $R_3$ radical is attached to the benzylic position only and/or it can also form a cyclic system by joining the C-2 position of phenyl. Preferred examples of $R_3$ are: -H, -OH, -$CH_3$ or a chain attached to the C-2 of phenyl: -CH=CH-CH=, -$CH_2$-O-, -$CH_2$-NH, -$CH_2$-S, -CH=CH-N=, =CH-O-, =CH-NH-, =CH-S- Each one of radicals $R_4$ and $R_5$ is attached at any possible substitution position of the benzene ring (ortho, meta, para), and they are independently selected from a group consisting of: H, F (and Cl, Br, I), $NO_2$, $CF_3$, $OCH_3$, $OCF_3$, $SCH_3$, (C1-C3)-alkyl, etc. Other $R_4$ and $R_5$ are possible, e.g.: -CN, $COCH_3$, COH, OH, $NH_2$, COOMe, COOEt.

Viral infection. A549 expressing hACE2 (A549.hACE2) were generated by transduction of parental A549 cells (ATCC CRM-CCL-185) with a retroviral vector expressing human ACE2 and a selection marker that confers resistance to blasticidin. pCMV3-hACE2 was obtained from Sino Biologicals S.L. (Beijing, China). SARS-CoV-2 (Coronaviridae; Orthocoronavirinae; Betacoronavirus; Sarbecovirus; strain NL/2020) was kindly provided by Dr. R. Molenkamp, Erasmus University Medical Center Rotterdam and was propagated in Vero-E6 cells as previously described [1](1). Virus stocks were titrated using tissue culture infectious dose 50 per ml (TCID50/ml) determination 2 in Vero-E6 cells and using immunofluorescence microscopy using an antibody against SARS-CoV-2 N protein (Genetex HL344) by end-point dilution and infection foci counting in A549-ACE2 cells. A549.hACE2 cells ($10^5$ cells/well) were seeded onto 12-well plates in DMEM/10%FCS. The day after, dilutions of the candidate compounds were prepared to obtain a 2X solution (50 µM; 10 µM for Remdesivir) in DMEM/2% FCS. Remdesivir (5 µM, final conc.) was used as a positive control for infection inhibition. On the other hand, a SARS-CoV-2 virus stock (NL/2020; alpha variant) was diluted in DMEM-2%FCS to achieve a multiplicity of infection of 1 pfu/cell. In addition to MOCK-treated samples (only media), comparable volumes of heat-inactivated virus stocks (30 min at 60 ºC) were used as a negative control for potential interference of VeroE6-derived factors during virus stock production. Equal volumes of diluted 2X compounds (50 µM; final concentration 25 µM) and virus stock were mixed prior to inoculation of target cells. Cells were incubated for 24 h at 5% $CO_2$, 37 ºC and 95% humidity. Media was removed and total cell extracts were produced by direct addition of 250 µl of trizol per well.

Ethics Statement: All the experiments involving virus infection of cell cultures were performed by the Antiviral Screening Platform in authorized biosafety level 3 (BSL-3) facilities of the Centro Nacional de Biotecnología (CNB; Madrid, Spain), following international regulations and under the CSIC Ethics Committee supervision.

### Supplemental Tables

**Table ST1. Average free-binding energies during the last 10ns and during the residence time of molecular dynamics and the residence time of candidate ORF9b homodimerization surface-binding hit compounds.**

| Ligand | $\Delta G_{ave}$ (10 ns) (kcal/mol) | $\Delta G_{ave}$ (RT) (kcal/mol) | RESIDENCE TIME (ns) |
|---|---|---|---|
| EN300 | -29,00 | -27,95 | 60 |
| Z118 | -27,15 | -27,15 | 10 |
| Y600 | -38,53 | -38,53 | 10 |
| Z268 | -37,20 | -34,65 | 110 |
| Y203 | -33,44 | -32,32 | 340 |
| K603 | -32,98 | -33,89 | 190 |
| Z166 (1) | -33,96 | -33,26 | 280 |
| Z166 (2) | -26,16 | -25,75 | 40 |
| Z158 | -24,33 | -23,78 | 110 |
| Z285 (1) | -40,30 | -40,3 | 10 |
| Z285 (2) | -45,94 | -44,06 | 30 |
| Z112 | -44,06 | -45,81 | 190 |

**Table ST2. Average free-binding energies during the last 10ns and during the residence time of molecular dynamics and the residence time of EN-300-similar compounds on the ORF9b homodimerization surface.**

| LIGAND | $\Delta G_{ave}$ (10 ns) (kcal/mol) | $\Delta G_{ave}$ (RT) (kcal/mol) | Residence Time (ns) |
|---|---|---|---|
| Compound 2 | -31,95 | -32,34 | 80 |
| Compound 4 | -26,12 | -25,08 | 200 |
| Compound 5 | -21,30 | -23,15 | 120 |

**Table ST3. Major interactions between ORF9b residues and EN300-series ORF9b homodimerization inhibitors**

**EN300** *kcal/mol

| Residue (position) | Total_ΔG* | St.Dev. | VdW | Elec | ΔGsolv* |
|---|---|---|---|---|---|
| **V15** | -1,3 | 0,3 | -0,7 | -0,4 | -0,2 |
| **L46** | -2,6 | 0,6 | -1,4 | -0,3 | -0,9 |
| **P51** | -2,4 | 0,8 | -1,1 | -0,5 | -0,8 |
| **L52** | -6,4 | 1,3 | -2,4 | -2,7 | -1,3 |
| **S53** | -1,4 | 0,3 | -0,9 | 0,1 | -0,6 |
| **L54** | -2,2 | 0,6 | -1,2 | -0,2 | -0,8 |
| **V76** | -2,3 | 0,5 | -1,1 | -1,0 | -0,3 |
| **Q77** | -4,7 | 1,2 | -1,3 | -2,9 | -0,5 |
| **M78** | -4,3 | 0,7 | -2,2 | -1,0 | -1,1 |
| **K97** | -1,9 | 1,7 | -0,9 | 0,1 | -1,1 |

**Compound 2** *kcal/mol

| Residue (position) | Total_ΔG* | St.Dev. | VdW | Elec | ΔGsolv* |
|---|---|---|---|---|---|
| **V15** | -1,0 | 0,5 | -0,6 | 0,2 | -0,6 |
| **L46** | -1,3 | 0,6 | -0,7 | 0,0 | -0,5 |
| **L52** | -2,7 | 1,2 | -1,4 | -0,1 | -1,2 |
| **S53** | -2,0 | 1,6 | -1,0 | -0,9 | -0,2 |
| **L54** | -2,7 | 2,1 | -1,3 | -0,6 | -0,7 |
| **A75** | -1,8 | 1,0 | -0,8 | -0,4 | -0,5 |
| **V76** | -4,6 | 1,7 | -1,6 | -2,2 | -0,8 |
| **Q77** | -1,2 | 1,2 | -0,7 | -0,2 | -0,2 |
| **M78** | -1,7 | 0,8 | -0,9 | -0,1 | -0,7 |
| **K97** | -2,6 | 2,0 | -0,8 | -3,1 | 1,2 |

**Compound 4** *kcal/mol

| Residue (position) | Total_ΔG* | St.Dev. | VdW | Elec | ΔGsolv* |
|---|---|---|---|---|---|
| **V15** | -1,1 | 0,4 | -0,6 | 0,0 | -0,6 |
| **L46** | -2,1 | 0,6 | -1,1 | 0,0 | -1,0 |
| **L48** | -1,3 | 0,7 | -0,8 | 0,1 | -0,7 |
| **P51** | -3,5 | 0,7 | -1,7 | -1,1 | -0,7 |
| **L52** | -6,2 | 1,1 | -3,1 | -1,4 | -1,7 |
| **S53** | -1,1 | 0,3 | -0,8 | 0,3 | -0,6 |
| **L54** | -1,7 | 0,5 | -0,9 | -0,1 | -0,7 |
| **V76** | -2,9 | 0,7 | -1,5 | -0,1 | -1,3 |
| **M78** | -3,1 | 0,8 | -1,6 | -0,6 | -0,9 |

#### Supplemental Figure

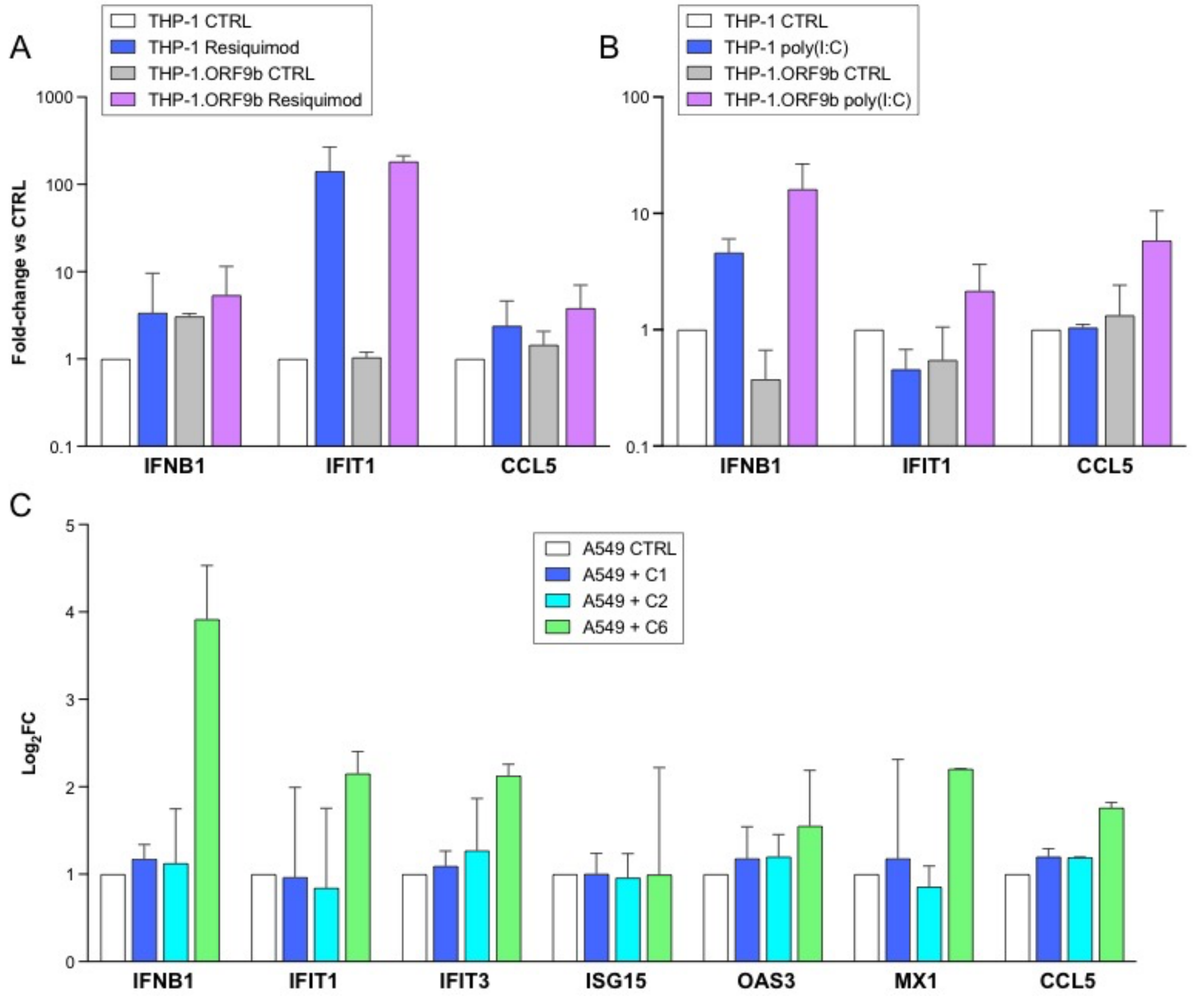


**Figure SF1. A, B**. ORF9b fails to antagonize the induction of ISGs by Resiquimod (**A**) or poly(I:C) (**B**). Wild-type THP-1 cells and THP-1 with stable integration and constitutive expression of ORF9b (THP-1.ORF9b) were exposed to Resiquimod (20 μM) or solvent (CTRL) for 4 h (A) or transfected with poly(I:C) (1 μg/mL) or vehicle (CTRL) for 4 h, and transcript levels of the indicated genes quantified by RT-PCR with TaqMan assays. Shown are fold-change values relative to the corresponding control conditions. All experiments were performed in triplicate. **C**. Effect of ORF9b dimerization inhibitors, Compounds 1, 2 or 6 (C1, C2, C6) on the basal expression levels of ISGs in A549.hACE2 cells in the absence of SARS-CoV-2 viral infection. Cells were exposed to 5 μM of each compound for 2 h, and transcript levels of the indicated genes quantified by RT-PCR with TaqMan assays.
